# Expression and Characterization of SARS-CoV-2 Spike Protein in *Thermothelomyces heterothallica* C1

**DOI:** 10.1101/2025.05.01.651343

**Authors:** Yakir Ophir, Justin Wong, Katherine R. Haddad, Anne Huuskonen, Anindya Karmaker, Varun Gore, Seongwon Jung, Armin Oloumi, Yiyun Liu, Jingxin Fu, Libo Zhang, Peishan Huang, Shiaki Arnett Minami, Shruthi Satya Garimella, Anugraha Thyagatur, Paulo A. Zaini, Marika Vitikainen, Ronen Tchelet, Noelia Valbuena, Thomas R. Fuerst, Emrullah Korkmaz, Louis D. Falo, Stephen C. Balmert, Saniya Mahendiratta, Mark Emalfarb, Priya S Shah, Justin Siegel, Abhaya M. Dandekar, Xi Chen, Carlito Lebrilla, Roland Faller, Markku Saloheimo, Karen A. McDonald, Somen Nandi

## Abstract

The COVID-19 pandemic demonstrated a pressing need for rapid, adaptive, and scalable manufacturing of vaccines and reagents. With the transition into an endemic disease and rising threats of other emerging pandemics, production of these biologicals requires a stable and sustainable supply chain and accessible distribution methods. In this study, we demonstrate the strength of an engineered filamentous fungal platform, *Thermothelomyces heterothallica* C1, for high volumetric productivity of the full-length spike glycoprotein. Spike protein produced in this system is highly thermostable and immunization of mice with spike made in C1 or mammalian platforms resulted in a similar humoral response. Additionally, it was shown that the native N-glycan profile can be redecorated with complex sialylated structures, if necessary, resulting in a more human-like glycan profile, without impacting binding characteristics as shown experimentally and in simulations. Through extensive physicochemical analysis, the C1 produced spike performs similarly to spike proteins produced in other commercially available systems. The data presented is evidence that C1 can be a strong platform for production of complex glycosylated recombinant proteins such as subunit antigen vaccines.

## Introduction

COVID-19 caused by the Severe Acute Respiratory Syndrome Coronavirus-2 (SARS-CoV-2) has over 770 million confirmed cases, with more than 7 million deaths as of March 2024 [1]. In the first year of vaccine availability (December 2020 – December 2021), it is estimated that 14.4 million deaths were prevented through vaccination [2]. The success of these vaccines, however, is not distributed equitably. As of November 2023, over 13.5 billion doses of vaccines have been administered globally [1], but there is a stark gap in the vaccination rates between high- and low-income countries [3]. As COVID-19 transitions from an acute health emergency into an endemic disease, the burden of procuring and distributing vaccines will now be placed on primary health care services instead of international organizations like the COVID-19 Vaccine Delivery Partnership. With this transition, there is a risk that disparities between high- and low-income countries will widen, as healthcare spending in low-income countries will have to increase by more than 50% to achieve a goal of 70% vaccination coverage, compared to a spending increase of just 0.8% for high-income countries [3].

Operational challenges such as inadequate cold chains, lack of transportation equipment, unpredictable supply chains, vaccine stability, and administrative logistics are all factors that contribute to the financial and logistical burdens that increase the cost and access of vaccines. In addition, SARS-CoV-2 variants of concern (VOCs) [4] have emerged that reduce the efficacy of previous vaccinations [5], creating a need to generate updated vaccine boosters on a regular basis. Consequently, a vaccine platform that is rapid, inexpensive and flexible will be needed to ensure equitable access to efficacious vaccinations for COVID-19 and other illnesses.

As of December 2022, several platforms have received approval for use as COVID-19 vaccines by at least one regulatory body globally, including nucleic acids (DNA/RNA), non-replicating viral vectors, inactivated virus, virus like particles, and protein subunit vaccines [6]. The platforms each have their own advantages and disadvantages as summarized by Chakraborty et al. [7]. Among these platforms, key advantages of recombinant protein subunit vaccines over other methods include ease of production, storage, transportation, and distribution, often making them more accessible and affordable [8]. In addition, the recombinant proteins used for vaccination can also be used for serological testing and diagnostic assays.

The majority of COVID-19 vaccines and vaccine candidates use the SARS-CoV-2 spike protein [9], a major structural glycoprotein found as a homotrimer on the surface of the viral envelope [10], as an antigenic target [11, 12]. Each spike protein monomer contains an S1 domain that is responsible for binding to the angiotensin-converting enzyme 2 (ACE2) cell receptor and an S2 subunit that is critical in mediating the fusion of the viral and host cell membranes [11]. Of the protein subunit vaccines approved for use [13] or in development [14], the predominant antigen of choice has been a region found in the spike S1 domain known as the receptor binding domain (RBD), responsible for the interaction of ACE2 and spike protein. Many platforms have been successful in producing RBD antigens, including mammalian, insect, plant, yeast, and bacterial systems, while the production of full-length spike protein has proven to be more difficult due to size and molecular complexity [15, 16, 17]. Studies have demonstrated that antigens covering regions outside of the RBD can produce increased levels of neutralizing antibodies in patients [18, 19], thus a vaccine that utilizes antigenic epitopes found along the entire spike protein may be beneficial for immunization.

Here, we present the production of the full-length spike protein (C1-Spike) in C1, a genetically modified thermophilic filamentous fungus (*Thermothelomyces heterothallica*) clonal cell line that has been optimized for glycoprotein expression [20, 21]. The functionality of C1-Spike is examined using enzyme-linked immunosorbent assay (ELISA) and biolayer interferometry (BLI), while protein structure and binding interactions are probed using circular dichroism (CD), liquid chromatography-tandem mass spectrometry (LC-MS/MS), and steered molecular dynamics (SMD) simulations. In addition, it is demonstrated that the N-glycan profile of C1-Spike can be modified to contain mostly monosialylated glycans that is more representative of human glycan profiles. Furthermore, immunogenicity of C1-Spike is explored through murine studies. Based on these data, C1-Spike can potentially be used as an antigen for vaccine candidates. To the best of our knowledge, this is the first report of successful production of full-length spike protein in filamentous fungi.

## Materials and Methods

### Construction of Full-Length Spike C1 Strain

The construction of a C1 fungal strain expressing the full length spike protein was done as previously published for SARS-CoV-2 RBD production [13], with modification as follows. A DNA coding sequence of SARS-CoV-2 strain Wuhan-Hu-1 surface glycoprotein residues 16-1209 (GenBank QHD43416.1) with mutated S1/S2 furin cleavage site (682-685 RRAR to GSAS) and double proline stabilization (986-987 KV to PP), a foldon trimerization domain with Gly-Ser-linker dipeptide (GSGYIPEAPRDGQAYVRKDGEWVLLSTFL) after residue 1209, followed by a carboxy-terminal tetrapeptide C-tag (EPEA) was designed and synthesized by Genscript (Piscataway, NJ, USA). The construct was inserted into the PacI restriction site of pMYT1055, the C1-cell expression vector plasmid, using seamless cloning and confirmed by DNA sequencing. Strain generation of C1 cells expressing C1-Spike is outlined in Supplemental Figure 1, following methods described in [13].

**Supplemental Figure 1.**
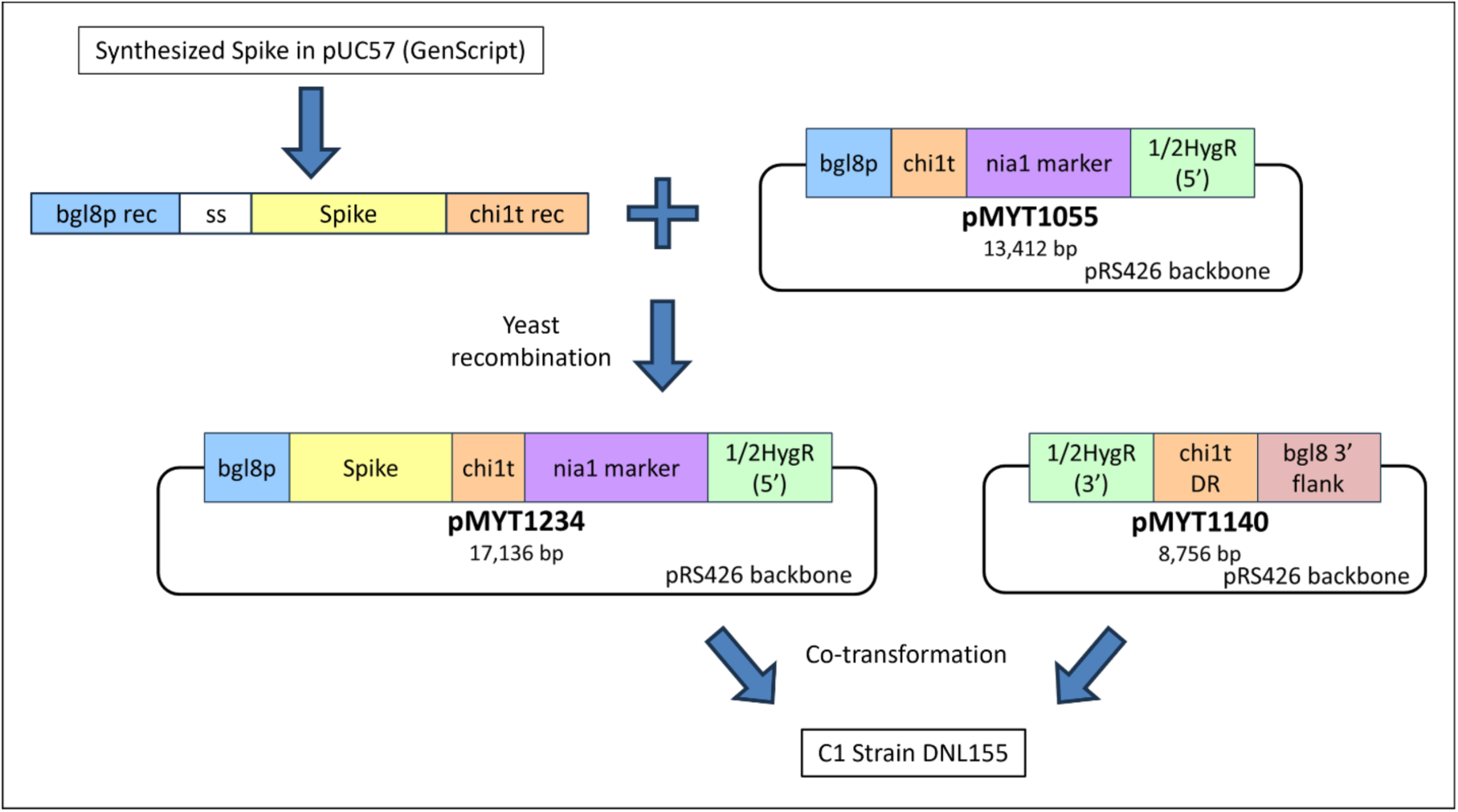
Flow chart of the construction of the Spike-C-tag expression plasmid pMYT1234. Plasmid numbers and size as base pairs (bp) indicated. Abbreviations: bgl8p, bgl8 promoter region; bgl8p rec, 40 bp recombination sequence to bgl8p; bgl8 3’ flank, 3’ flank region of bgl8; ss, cellobiohydrolase 1 signal sequence; Spike, Spike protein encoding amino acids 16-1209 of the Spike protein from SARS-CoV-2 including furin cleavage site, double proline stabilization, foldon trimerization domain, and C-tag; chi1t, terminator; chi1t rec, 40 bp recombination sequence to chi1t; chi1t DR; direct repeat fragment of chi1t; nia1 marker, nitrate reductase marker for transformant selection; HygR (5’ and 3’ halves), hygromycin resistance marker for transformant selection.

### Fermentation

Seed cultures were prepared by thawing 1 mL of a frozen glycerol stock and adding it to a 250 mL shake flask containing 49 mL of complete media as described in [13]. This culture was incubated for 2 days at 37°C with shaking at 250 rpm. After 2 days, the culture was split into two 4 L baffled shake flasks, where 25 mL of culture was added to 300 mL of complete media. These flasks were incubated at 37°C with shaking at 250 rpm for 24 h.

C1-Spike was produced in a BioFlo3000 5 L bioreactor equipped with two Rushton impellers (New Brunswick, NJ, USA) using a fermentation process as previously reported [13], with slight modification to the starting volumes, temperature setpoints, dissolved oxygen (DO) agitation limits, nutrient feed rates, and the addition of media exchanges during the fermentation. In brief, 300 mL of seed culture was added to 3 L of autoclave-sterilized high concentration yeast extract media as prepared in [13]. An additional 500 mL of fresh culture media was added at 22 h elapsed fermentation time (EFT) to submerge the second impeller. The fermentation was initiated at 35°C; with the temperature reduced to 30°C at 20 h EFT and 25°C at 73.7 h EFT, which was maintained for the remainder of the fermentation. The DO setpoint of 20% air saturation was maintained by agitation control at 150-800 rpm and an air flow rate set to 4 L/min. During the fermentation, the air flow rate was reduced to 2 L/min due to exhaust filter fouling. The exhaust filter was replaced at approximately 70 h EFT, and the air flow rate recovered to 4 L/min. Despite this decrease in air flow rate, the DO was well controlled at 20%. A nutrient feed with the same composition as reported in [13] was started at a rate of 7 mL/hr at 29 h EFT and reduced to 5 mL/hr at 77.6 h EFT to maintain a packed cell volume ratio of 25% per sample volume. At 72.2 h EFT, 1 L of culture broth was withdrawn and exchanged with an equal volume of fresh media. This was repeated at 94 h EFT by exchanging a volume of 0.5 L. During the media exchanges, the feed was paused for a duration of 4 h before restarting.

### Media Clarification and Concentration

Broth from the fermentation harvest at 115 h EFT was centrifuged for 25 min at 20,000 x g and 4°C using an Avanti JXN-26 Centrifuge (Beckman Coulter, IN, USA). The supernatant was collected and filtered through a Supracap 50 depth filter with a 0.5-15 μm retention rating (Pall, NY, USA). The clarified solution was loaded onto an Äkta Flux tangential flow filtration (TFF) system (Cytiva, MA, USA), where it was buffer exchanged into 1x phosphate-buffered saline (PBS), pH 7.2, and concentrated eight-fold on a Pellicon XL50 with a Biomax 30 kDa membrane (MilliporeSigma, MA, USA). A recirculation rate of 50 mL/min was maintained for the duration of the process, and the retentate valve was opened as needed during the concentration step to maintain a transmembrane pressure of 35 psig. Concentrated TFF retentate was stored at -20°C until purification.

### Protein Purification

Purification was performed on an Äkta Pure (Cytiva, MA, USA) fast protein liquid chromatography system using an XK16 column, with an inner column diameter of 1.6 cm (Cytiva, MA, USA) packed with 5 mL of NGL COVID-19 Spike Protein Affinity Resin (Repligen, MA, USA). Samples for purification were centrifuged at 20,000 x g and filtered through a 0.45 μm filter before column loading. The column was equilibrated with PBS, pH 7.2 and then loaded with the concentrated TFF retentate. The column was washed with 10 column volumes (CV) of 0.51 M sodium chloride (NaCl), pH 7.0, at a flow rate of 1 mL/min and bound proteins were eluted with a single-step gradient using Pierce Gentle Ag/Ab Elution Buffer, pH 6.6 (Thermo Fisher Scientific, MA, USA) for 5 CV at a flow rate of 0.5 mL/min. Eluted samples were dialyzed against a 0.16 M NaCl, pH 7.0 solution overnight at 4°C in 20 kDa Slide-A-Lyzer G2 Dialysis Cassettes (Thermo Fisher Scientific, MA, USA). Dialysis samples were then concentrated ten-fold on 100 kDa Amicon Ultra 0.5 Centrifugal units (MilliporeSigma, MA, USA) before analysis and storage.

The purity of eluted sample was determined using gel densitometry methods on Image Lab software v6.1 (Bio-Rad Laboratories, CA, USA) and concentration was determined by absorbance at 280 nm with a SpectraDrop Micro-Volume Microplate and SpectraMax M4 Microplate Reader (Molecular Devices, CA, USA). An extinction coefficient of 135,720 M^-1^cm^-1^ as predicted by ExPASy ProtParam [22] was used to calculate the concentration of C1-Spike from the absorbance reading.

### Protein Electrophoresis and Immunoblotting

Sodium dodecyl sulfate polyacrylamide gel electrophoresis (SDS-PAGE) and Western blot were used to evaluate the production of C1-Spike during fermentation, the purity of C1-Spike after downstream processing, and the integrity of the protein during processing steps. Samples were heated to 95°C for 5 min in 4x Laemmli buffer (Bio-Rad Laboratories, CA, USA) containing 50 mM of dithiothreitol (DTT). Aliquots of 30 μL of the heat-treated samples were loaded onto precast 4-20% SDS-Tris HCl polyacrylamide gels (Bio-Rad Laboratories, CA, USA), with 10 μL of Precision Plus Protein All Blue Standards (Bio-Rad Laboratories, CA, USA) and 10 μL of Precision Plus Protein Unstained Standards (Bio-Rad Laboratories, CA, USA) loaded into separate lanes. SDS-PAGE was performed in a Mini-PROTEAN Tetra Vertical Electrophoresis Cell (Bio-Rad Laboratories, CA, USA) powered by a PowerPac Basic (Bio-Rad Laboratories, CA, USA) power supply at 90 V for 90 min using Tris/Glycine/SDS running buffer (Bio-Rad Laboratories, CA, USA). After electrophoresis, gels were either stained with Coomassie Brilliant Blue R-250 Staining Solution (Bio-Rad Laboratories, CA, USA) or utilized for Western blot analysis. Stained gels were used for purity confirmation by densitometry using ImageLab v6.1 (Bio-Rad Laboratories, CA, USA).

Samples for Western blotting were transferred onto a 0.2 μm nitrocellulose membrane (Bio-Rad Laboratories, CA, USA) using the Trans-Blot Turbo Transfer System (Bio-Rad Laboratories, CA, USA). The membrane was then blocked with 1% w/v Casein in PBS (Bio-Rad Laboratories, CA, USA) for 1 h at room temperature. For Western blot detection, the membrane was probed overnight with a rabbit anti-spike protein (SARS-CoV-2) polyclonal antibody (eENZYME, MD, USA) at a concentration of 1 μg/mL in blocking buffer, followed by 1 h incubation with a goat anti-rabbit IgG (H+L), Mouse/Human ads-HRP antibody (SouthernBiotech, AL, USA) at a 1:1000 dilution in blocking buffer. Blots were developed using Clarity Western ECL substrate (Bio-Rad Laboratories, CA, USA) and imaged using the ChemiDoc System (Bio-Rad Laboratories, CA, USA).

### ELISA Quantification

High binding 96-well plates (Corning, MA, USA) were coated with crude fermentation supernatant or purified C1-Spike protein standard starting at 0.5 µg/mL and diluted two-fold in 0.16 M NaCl. Adsorption occurred at 4°C overnight and plates were allowed to reach room temperature (RT) before washing. Plates were washed three times in PBS plus 0.05% v/v Tween (PBST) and blocked for 1 h at RT in 1% w/v Casein in PBS (Bio-Rad Laboratories, CA, USA). C1-Spike was sequentially probed with the C1 produced anti-RBD 87G7 monoclonal antibody [23, 24] at 0.1 µg/mL and HRP conjugated anti-human IgG (SouthernBiotech, AL, USA) at 0.275 µg/mL, both diluted in blocking buffer for 1 h at RT. Reaction was developed with TMB/E substrate (MilliporeSigma, MA, USA) and stopped with 1N hydrochloric acid. Signal was read at 450 nm on a SpectraMax M4 Microplate Reader (Molecular Devices, CA, USA). Optical density was plotted against concentration, and data were fitted to a 4-parameter logistic model using SoftMax Pro Software v7.1 (Molecular Devices, CA, USA). Error bars represent the standard deviation of samples run in triplicate and average values were determined by readings on the linear region of the standard curve.

### Antibody Binding Characterization

Three standard curves were generated following the indirect ELISA format as described above. C1-Spike was serially diluted two-fold from 0.5 µg/mL in 0.16 M NaCl in duplicate. In-house C1 produced 87G7 [23, 24], HEK293 produced S2H97 [25], and HEK293 produced REGN10933 [26] were used to probe the antigen at a concentration of 0.1 µg/mL. Graphs of all three antibodies were overlaid.

### Biolayer Interferometry

To determine the kinetics of interactions between C1-Spike to its receptor protein ACE2, anti-Human IgG Fc Capture (AHC) biosensors (ForteBio, CA, USA) were used to immobilize ACE2 conjugated to fragment crystallizable region of IgG1 (ACE2-Fc) (AcroBiosystems, DE, USA) by immersing the AHC biosensors into kinetic buffer (Sartorius, Göttingen, Germany) containing 100 nM ACE2-Fc for 10 min. The kinetics of C1-Spike interacting with reported neutralizing antibody 87G7 [23] were determined by immobilizing the C1-produced 87G7 on AHC biosensors by immersing the biosensors into a kinetic buffer containing 21 nM of 87G7 for 10 min. The Octet RED384 (ForteBio, CA, USA) was used to obtain response measurements for protein association and dissociation. The AHC sensors with either ACE2-Fc or 87G7 were then dipped into two-fold dilutions of either C1-Spike or glycan modified C1-Spike from 125 to 3.91 nM in kinetic buffer. The dissociation phase was also conducted in kinetic buffer. Data were collected for 90 s for baseline, 1200 s for association, and 1200 s for dissociation; experiments were conducted at 37°C. AHC biosensors were regenerated using 10mM Glycine-HCl, pH 1.5.

ForteBio Data Analysis Software v8.1 (ForteBio, CA, USA) was used for data pre-processing and analysis. Raw data were reference well subtracted, the y-axes were aligned to baseline, interstep correction was applied for alignment to dissociation, and Savitzsky-Golay filtering was used for smoothing. A 1:1 binding model with a global fit and R_max_ unlinked by sensor was used to determine kinetic parameters.

### Circular Dichroism

Samples were analyzed by the Protein Structure and Dynamics Core at the University of California, Davis as published before [27]. Briefly, 150 µg of protein in 300 µL PBS was provided, and singular spectral data were obtained over a 200-250 nm wavelength range on a JASCO J-715 CD spectrometer (JASCO, MD, USA). Results were analyzed using BeStSel webserver [28, 29]. Running buffer spectra were subtracted from sample signals and a scale factor of 0.1 was used across samples to determine the percentages of secondary structures. C1-Spike structure was compared to spike proteins produced in *Spodoptera frugiperda* (Sf9) or Chinese hamster ovary (CHO) cells. S9-Spike and CHO-Spike contain the same 986-987 KV to PP stabilization as C1-Spike, while CHO-6P-Spike refers to the six-proline stabilization termed HexaPro [15].

### Temperature Stability

The stability of C1-Spike was evaluated over 7 days at 4°C, stored in the refrigerator (VWR, PA, USA) or at 37°C, stored in an EchoTherm IN-35 incubator (Torey Pines Scientific, CA, USA). A 50 μL sample was taken each day and stored at -80°C until further analysis by ELISA, as described above. Samples run on ELISA were normalized to Day 0 initial sample and a *p*-value was calculated using a one-way ANOVA.

### Lyophilization

C1-Spike was lyophilized on a FreeZone 4.5 L freeze dry system (Labconco, MO, USA). Before the sample was loaded, the lyophilizer was cooled below -40°C and maintained a vacuum of less than 0.133 mBar. A 1.2 mL sample of purified C1-Spike in a 7 mL borosilicate scintillation vial was flash frozen in liquid nitrogen and loaded onto the system for 48 hours. The dried sample was then reconstituted in 1.2 mL of deionized water and was gently agitated by hand every 2 min for 10 min at room temperature.

Samples of C1-Spike before and after lyophilization were run on ELISA, as described above. ELISA data were normalized to before treatment values and a *p*-value of lyophilization treatment was calculated using Student’s two-tailed t-test.

### Glycoproteomic Analysis

Details of protein digestion for glycopeptide analysis have been described previously [30, 31]. Briefly, buffer exchange was performed using a 3 kDa spin column (Merck Millipore, MA, USA) to remove salts and dilute recombinant proteins with 50 µL of 50 mM ammonium bicarbonate solution, then the proteins were reduced with 2 μL of 550 mM DTT and alkylated with 4 μL of 450 mM iodoacetamide. The samples were divided into three fractions and incubated with trypsin, chymotrypsin, and α-Lytic protease at 37°C for 18 h. The resulting peptides and glycopeptides were dried using a miVac (SP Scientific, PA, USA) prior to mass spectrometry analysis.

The peptide and glycopeptide samples were reconstituted with nanopure water and directly characterized using UltiMate WPS-3000RS nanoLC 980 system coupled to the Nanospray Flex ion source of an Orbitrap Fusion Lumos Tribrid Mass Spectrometer system (Thermo Fisher Scientific, MA, USA). The analytes were separated on an Acclaim PepMap 100 C18 LC Column (3 μm, 0.075 mm × 150 mm, Thermo Fisher Scientific, MA, USA). A binary gradient was applied using 0.1% v/v formic acid in (A) water and (B) 80% acetonitrile: 0–5 min, 4–4% (B); 5–133 min, 4–32% (B); 133–152 min, 32%–48% (B); 152–155 min, 48–100% (B); 155–170 min, 100–100% (B); 170–171 min, 100–4% (B); 171–180 min, 4–4% (B). The instrument was run in data-dependent mode with 1.8 kV spray voltage, 275°C ion transfer capillary temperature, and the acquisition was performed with the full MS scanned from 700 to 2000 m/z in positive ionization mode. Stepped higher-energy C-trap dissociation at 30 ± 10% was applied to obtain tandem MS/MS spectra with m/z values starting from 120.

Glycopeptide fragmentation spectra were annotated using Byonic software (Protein Metrics, CA, USA) against the protein sequences. Common modifications, including cysteine carbamidomethyl, methionine oxidation, asparagine deamidation and glutamine deamidation were assigned. A published in-house N-glycan library [30] was utilized for the glycopeptide identification, and relative abundance values were calculated based on the glycopeptide precursor peak areas.

### N-Glycan Modification

*In vitro* glycoprotein N-glycan processing has recently been described [32]. This process, with slight modification as follows, was used to edit the N-glycoform of C1-Spike protein. One-pot six-enzyme reactions were carried out in a 0.5 mL microcentrifuge tube by incubating 0.33 mg C1-Spike protein, 2 mM UDP-GlcNAc, 2 mM UDP-Gal, 5% w/w EfMan-I-His_6_, and 5% w/w MBP-Δ28hGnT-I-His_6_, 5% w/w Δ24Bt3994-His_6_ and 5% w/w Δ18Bt1769-His_6_, 5% w/w MBP-Δ27hGnT-II-His_6_, 10% w/w MBP-Δ128Bβ4GalT1-His_6_ in 100 mM Tris-HCl, pH 7.5 containing 2 mM MgCl_2_, 2 mM CaCl_2_, and 2 mM MnCl_2_ at 30°C for 2 h. Then, 5 mM CMP-Neu5Ac and 5% w/w MBP-Δ89hST6GAL-I-His_6_ were added to the reaction mixture followed by incubation at 30°C overnight before purification and glycoproteomic analysis performed as described above.

### Computational all-atom C1-Spike and ACE2 Complex Model Building

Models were retrieved from the work of Casalino et al. [33], with additions of glycans before and after modification as identified by glycoproteomic analysis via CHARMM-GUI webserver [34]. Simulation configuration employed the all-atom Amber ff19SB force field for amino acids [35] and GLYCAM_06j for glycosylation [36]. Models were solvated in TIP3P water, neutralized with ions, and supplemented with 0.155 mM NaCl. Energy minimization was conducted using the steepest descent method. During the equilibration phase, a constant temperature of 37°C was maintained with the use of a velocity-rescale thermostat [37], which had relaxation times of 1 ps. Equilibration continued under the Berendsen thermostat [38] for 2 ns at a temperature of 37°C and pressure of 1 bar, with respective relaxation times of 1 ps and 12 ps. Further equilibration for 20 ns was conducted with a Parrinello-Rahman isotropic barostat [39], also featuring relaxation times of 1 ps and 12 ps. A cutoff scheme was implemented for nonbonded interactions, which were truncated at a distance cutoff of 1.1 nm, and Particle-mesh Ewald electrostatics were used with a cutoff of 1.1 nm.

Pulling simulations were conducted using GROMACS v2020.4 [40, 41]. The reaction coordinate aimed to disrupt key hydrogen bonds, perpendicular to the receptor binding domain and ACE2 interface, guided by literature [42, 43]. A spring constant of 1350 kJ/mol/nm² and a pull rate of 0.001 nm/ps were used under a Nose-Hoover thermostat [44, 45] and Parrinello-Rahman barostat at 1 atm. Two separate pulls with unique random seed velocities ensured reliability, pulling until at least a 10 nm separation was achieved between ACE2 and the C1-Spike trimer.

### Immunization of Animals

Female C57BL/6 mice (Jackson Laboratory, ME, USA) were housed under pathogen-free conditions at the University of Pittsburgh School of Medicine. Experiments were conducted with 6–8-week-old mice in accordance with Institutional Animal Care and Use Committee guidelines. C1-Spike without glycan modification was purified and stored frozen prior to formulation for injection. Mice (n = 5 for each group) were immunized by intramuscular injections of 5 µg C1-Spike or HEK-Spike with 50 µg Alhydrogel (InvivoGen, CA, USA) in 50 µL PBS into the hindlimb gastrocnemius muscle and boosted similarly 14 days later. For total IgG antibody measurements and neutralization assays, blood was collected from the saphenous vein 6 weeks after primary immunization, and serum was obtained by centrifugation at 10,000 rpm for 10 min. The isolated serum was stored at -20°C until use in ELISAs and neutralization assays described below.

### Measurement of Binding Antibody Responses

Serum SARS-CoV-2 spike protein-specific total IgG antibody levels were measured by indirect ELISA, as previously described [46, 47]. High-binding plates (Corning, MA, USA) were coated with 1 μg/mL spike protein (Sino Biological, PA, USA) in PBS and incubated overnight at 4°C. Plates were washed with PBST three times and blocked with 1% normal goat serum (Jackson ImmunoResearch, PA, USA) in PBS for 1 h at 37°C. Three-fold serial dilutions of serum in blocking buffer were added to plates and incubated for 2 h at 37°C. After washing, plates were incubated for 1 h at 37°C with 75 ng/mL biotinylated goat anti-mouse IgG secondary antibody (Jackson ImmunoResearch, PA, USA) in blocking buffer, washed, and incubated for 30 min at 37°C with 1:1000 streptavidin-HRP (BD Biosciences, NJ, USA) diluted in blocking buffer. After washing three times, plates were developed with 1-Step Slow TMB substrate (Thermo Fisher Scientific, CA, USA) for 20 min at room temperature, and the reaction was quenched with 1 M sulfuric acid. Optical density at 450 nm was read with a SpectraMax iD5 Hybrid Multi-Mode Microplate Reader (Molecular Devices, CA, USA). Titers were fit using least squares regression with a nonlinear asymmetric five-parameter dose-response curve for reciprocal dilutions versus OD450 (GraphPad Prism v10), and endpoint titers were calculated by interpolation with a threshold equal to the mean + 3 SD of OD450 for naïve serum samples.

### Neutralization Assay

SARS-CoV-2-specific neutralizing antibodies in sera were measured as previously described [47]. HEK-293T-hACE2 cells (BEI Resources, VA, USA) were plated at 1.5 × 10^4^ cells/well in 50 μL media (DMEM with L-glutamine, sodium pyruvate, HEPES, and 10% FBS) in 96-well plates (Corning, MA, USA) coated with 0.01% poly-L-lysine (MilliporeSigma, MA, USA). The next day, serum samples were heat-inactivated at 56°C for 30 min and then diluted with media in 96-well round-bottom plates (Corning, MA, USA). Replication-deficient murine leukemia virus pseudotyped with the SARS-CoV-2 spike protein (GenBank QHD43416.1) and containing a firefly luciferase open reading frame as a reporter (MyBioSource, CA, USA) was added at 60 μL pseudovirus to 20 μL of three-fold serially diluted serum resulting in serum ranges from 1:20 to 1:131220. Serial dilutions of anti-Spike RBD neutralizing antibody (Sino Biological, PA, USA) were used to validate the assay. Pseudovirus was incubated with diluted serum, or neutralizing antibodies, for 2 h at 37°C, and then 70 μL/well transferred to cells. Polybrene (MilliporeSigma, MA, USA) was added with a final concentration of 5 μg/mL and a final volume of 150 μL/well. Cells were incubated at 37°C with 5% CO2 for 42 h, and then luciferase activity was measured using a ONE-Glo Luciferase Assay System (Promega, WI, USA). Lysed cells in ONE-Glo reagent were transferred to white plates (Corning, MA, USA) and luminescence read on a SpectraMax iD5 Microplate Reader (Molecular Devices, CA, USA) with 1 s integration. After subtracting background luminescence from control wells without pseudovirus, percent inhibition of pseudovirus infection was calculated by normalizing luminescence to that from control wells with cells and pseudovirus. Half maximal inhibitory dilution (ID50) neutralizing titers were determined by robust nonlinear regression using an inhibitor vs. normalized response, variable slope model, where % Inhibition = 100/(1+(ID50/Dilution) HillSlope), with ID50 > 0 and HillSlope < 0 constraints, and medium convergence criteria (GraphPad Prism v10).

## Results and Discussion

### C1-Spike Production

An expression plasmid for the C1-Spike was made by cloning the codon-optimized synthetic gene between the *bgl8* promoter and *chi1* terminator in a C1 expression vector (Supplementary Figure 1). The expression vector was transformed into the C1 strain DNL155 that has 14 deletions of protease genes [13, 21]. The expression cassette integrated as single copy into the native *bgl8* locus where high expression can be reached, and final production strains were shown by PCR to be genetically pure, i.e. devoid of any parental strain genetic material.

A selected C1-Spike production strain was cultivated in fed-batch fermentation for 115 hours. During the fermentation, C1-Spike secreted into the media was analyzed by SDS-PAGE, Western blot, and ELISA. A band at approximately 160 kDa corresponding to C1-Spike protein was detected starting at 54 h (Figure 1A, B lane 4) with increasing intensity over time (Figure 1A, B, lanes 5-9). Although the theoretical molecular weight (MW), excluding glycosylations, of C1-Spike is predicted to be 135 kDa by ExPASy ProtParam [22], the observed size of approximately 160 kDa is likely due to glycosylation [48].

**Figure 1.**
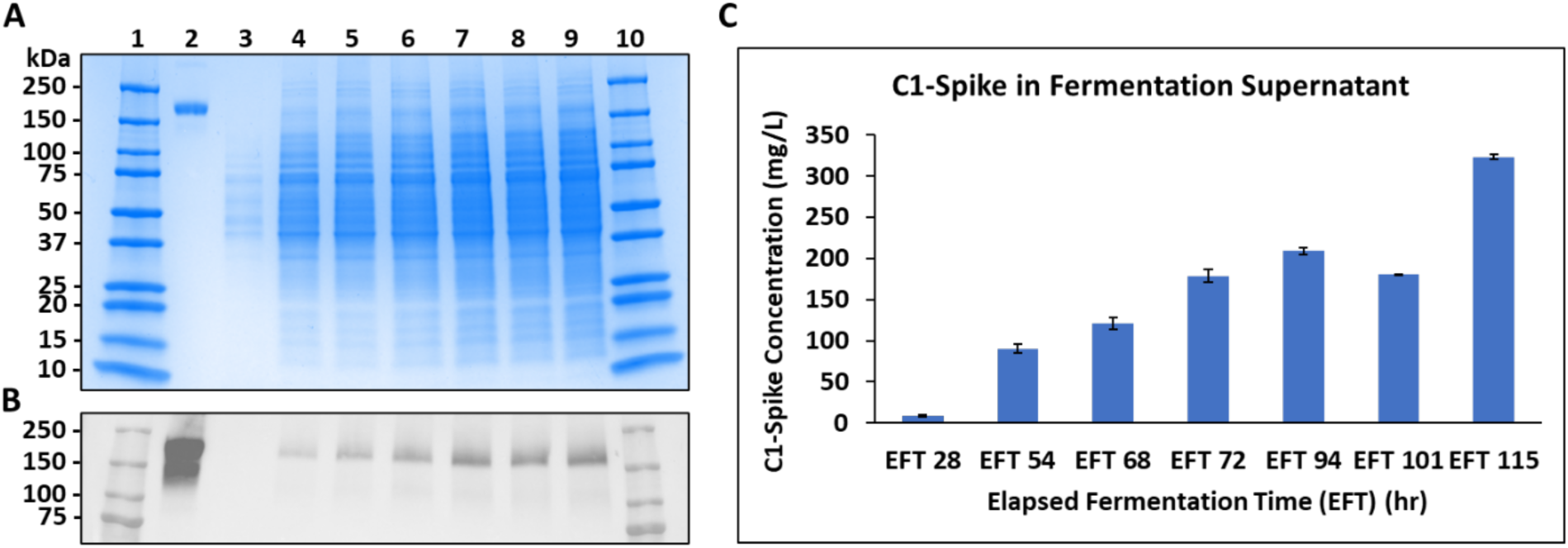
Time course of C1-Spike production during the C1 fermentation process. A) SDS-PAGE stained with Coomassie Blue. Lane 1, molecular weight marker (MWM); lane 2, reference CHO-Spike (BEI, NR-53937); lane 3, 28 h EFT (10x dilution); lane 4, 54 h EFT (10x dilution); lane 5, 68 h EFT (10x dilution); lane 6, 72 h EFT (10x dilution); lane 7, 94 h EFT (10x dilution); lane 8, 101 h EFT (10x dilution); lane 9, 115 h EFT (10x dilution); lane 10, MWM. B) Western blot of samples run in A. C) Indirect ELISA quantification of C1-Spike secreted in supernatant at various timepoints during fermentation. Error bars represent the standard error of averaged samples run in triplicate and averaged by those that lie on the linear region of the standard curve.

ELISA shows detection of C1-Spike beginning at 28 h EFT, with a marked increase in titer by 54 h (Figure 1C), as seen on Western blot (Figure 1B). C1-Spike titers continue to increase over time, with a slight increase in concentration between 72 h and 94 h EFT and a reduction in concentration between 94 h and 101 h EFT reflective of the dilutional impact of partially harvesting fermentation broth and replacing the volume with fresh media at 72.2 h and 94 h, respectively. These media exchanges allow earlier access to target protein which may prevent protein degradation and maintain culture productivity [49, 50]. C1-Spike fermentation was completely harvested at 115 h EFT, where production was approximately 320 mg/L as measured by ELISA (Figure 1C). Trends from ELISA indicate that the maximum titer of C1-Spike may not have been reached at the time of harvest, as this fermentation was not optimized for spike production, in contrast with previously reported production of RBD in a C1 system [13].

### Purification

Samples from the purification of C1-Spike protein from fermentation broth are visualized in Figure 2, where crude supernatant (Lane 2) is concentrated eight-fold by TFF (Lane 3) and purified on affinity resin (Lane 4). The loading mass of purified protein is determined using absorbance at 280 nm. A reference standard (Lane 5) was produced in CHO cells. The theoretical molecular weight of C1-Spike is 135 kDa, while the reference CHO spike has an expected molecular weight of 140 kDa, not including glycosylation. In Figure 2A, a Coomassie stained SDS-PAGE indicates minor host cell protein bands remaining in the purified elution, notably around 50 kDa and 75 kDa. Figure 2B shows a faint band at 100 kDa that could indicate degraded C1-Spike protein concentrated during the TFF process; however, these byproducts appear to be removed during affinity purification. A purity of approximately 88% was determined by gel densitometry of Lane 4 in Figure 2A. While this relatively high purity of C1-Spike was achieved through a one-step purification scheme, additional processing such as optimized washing conditions or secondary columns may be used to improve purity. The amino acid sequence of the purified protein was analyzed by LC-MS and the lack of any mutations were confirmed (Supplemental Figure 2).

**Figure 2.**
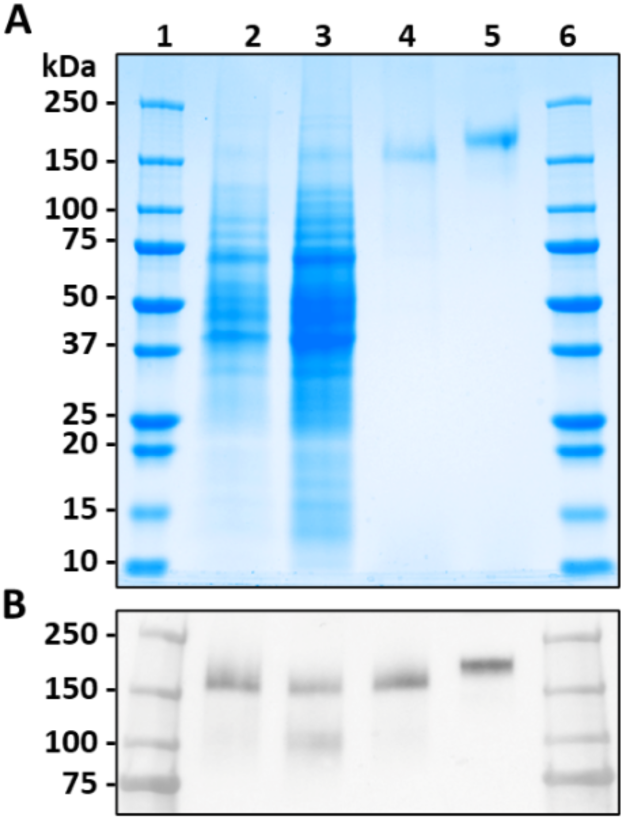
Purification processing for C1-Spike. A) SDS-PAGE stained with Coomassie Blue. Lanes 1 and 6, molecular weight marker (MWM); lane 2, 115 h EFT (20x dilution); lane 3, TFF retentate (30x dilution); lane 4, purified C1-Spike (3 µg); lane 5, reference CHO-Spike (BEI NR-53937) (3 µg). B) Western blot of samples in A, with 250 ng loading mass in lanes 4 and 5.

**Supplemental Figure 2.**
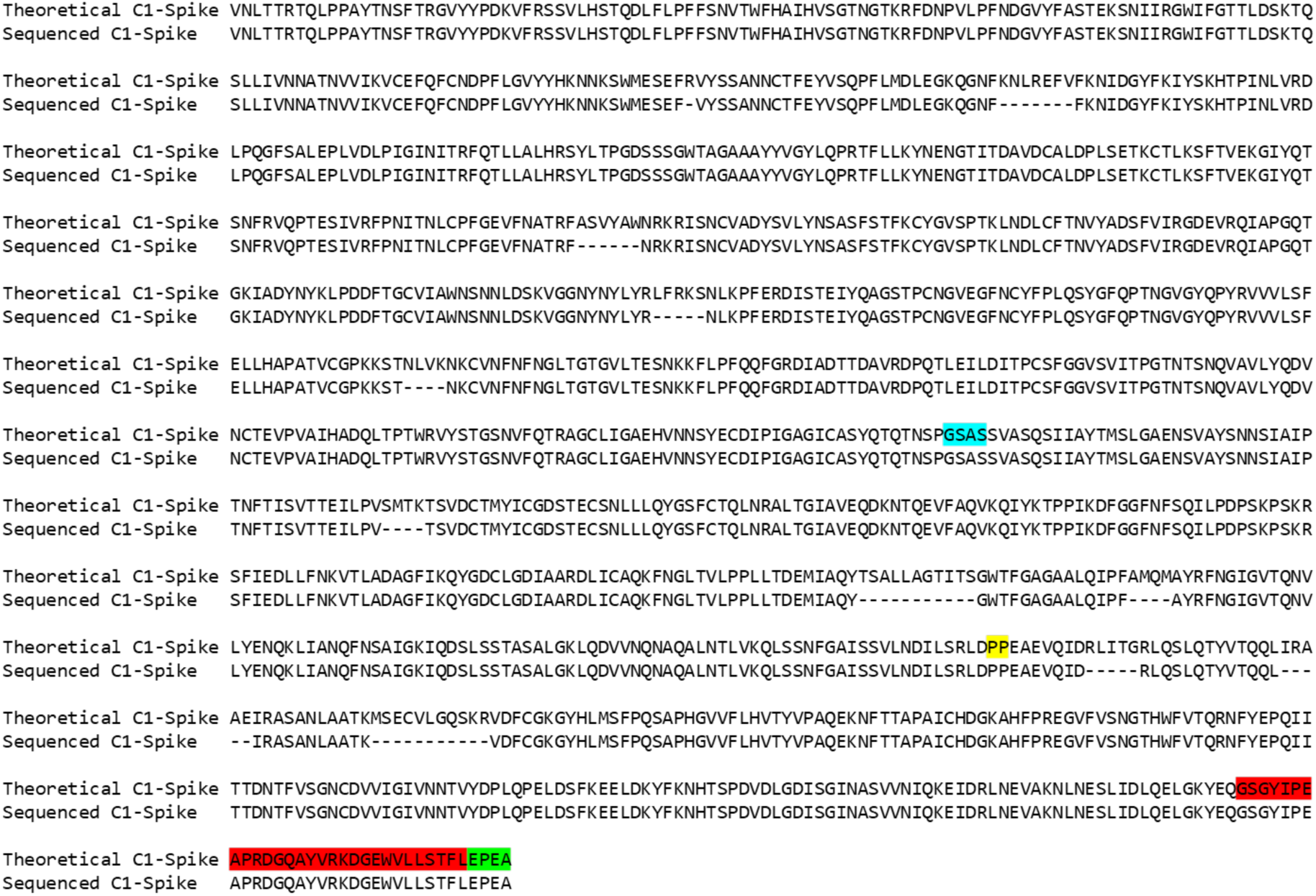
Amino acid sequence determination. A combined digestion of trypsin, chymotrypsin, and α-Lytic proteases of C1-Spike identified peptides through LC-MS with nearly 95% sequence coverage. Undetected amino acids are represented as dashes, and results indicate C1-Spike sequence matched with it’s native counterpart. Blue highlight denotes furin cleavage site; yellow, double proline mutation; red, foldon trimerization domain; green, C-tag.

### Binding Kinetics

Initial binding of C1-Spike to known antibodies was determined using ELISA, indicating that the antigen performed as previously reported [23, 24] (Supplemental Figure 3). Next, the binding kinetics between C1-Spike and ACE2 receptor or 87G7 antibody were measured by BLI (Table 1). The equilibrium dissociation constant (K_D_), association rate constant (k_on_), and dissociation rate constant (k_off_) of the interacting pairs can be found in Table 1. Binding curves are shown in Supplemental Figure 4. The K_D_ value of 48.8 nM for the ACE2 interaction with C1-Spike is consistent with the range of 6-133 nM observed by other groups [51]. N-glycan modifications of C1-Spike resulted in an order of magnitude reduction of K_D_ from 48.8 nM to 2.54 nM. However, this shift is considered to fall within the magnitude range of previously observed values of K_D_ for C1-Spike and ACE2 interactions. The K_D_ of C1-Spike and 87G7 could not be determined because there was negligible dissociation observed over 20 min with the conditions tested. Other groups [23] have reported dissociation at 30 min and report a binding affinity at sub nanomolar levels, consistent with the findings of this study.

**Table 1.** Binding constants for C1-Spike as determined by biolayer interferometry. Kinetic parameters were determined by fitting sensorgrams (Supplemental Figure 3) with the ForteBio Data Analysis Software v8.1 package.

| Ligand | Analyte | $K_D$ (M) | $K_D$ Error | $k_{on}$ (1/Ms) | $k_{on}$ Error | $k_{off}$ (1/s) | $k_{off}$ Error |
| --- | --- | --- | --- | --- | --- | --- | --- |
| ACE2-Fc | C1-Spike | 4.88E-10 | 1.48E-11 | 4.97E+04 | 1.34E+02 | 2.42E-05 | 7.33E-07 |
| 87G7 | C1 Spike | <1.0E-12 | 1.95E-11 | 8.05E+04 | 4.47E+02 | <1.0E-07 | N/A |
| ACE2-Fc | Glycan Modified C1-Spike | 2.54E-09 | 4.07E-11 | 2.78E+04 | 1.09E+02 | 7.05E-05 | 1.10E-06 |

| Ligand | Analyte | K<sub>D</sub> (M) | K<sub>D</sub> Error | k<sub>on</sub> (1/Ms) | k<sub>on</sub> Error | k<sub>off</sub> (1/s) | k<sub>off</sub> Error |
| --- | --- | --- | --- | --- | --- | --- | --- |
| ACE2-Fc | C1-Spike | 4.88E-10 | 1.48E-11 | 4.97E+04 | 1.34E+02 | 2.42E-05 | 7.33E-07 |
| 87G7 | C1-Spike | <1.0E-12 | 1.95E-11 | 8.05E+04 | 4.47E+02 | <1.0E-07 | N/A |
| ACE2-Fc | Glycan Modified C1-Spike | 4.80E-09 | 7.47E-11 | 2.60E+04 | 1.14E+02 | 1.25E-04 | 1.86E-06 |

**Supplemental Figure 3.**
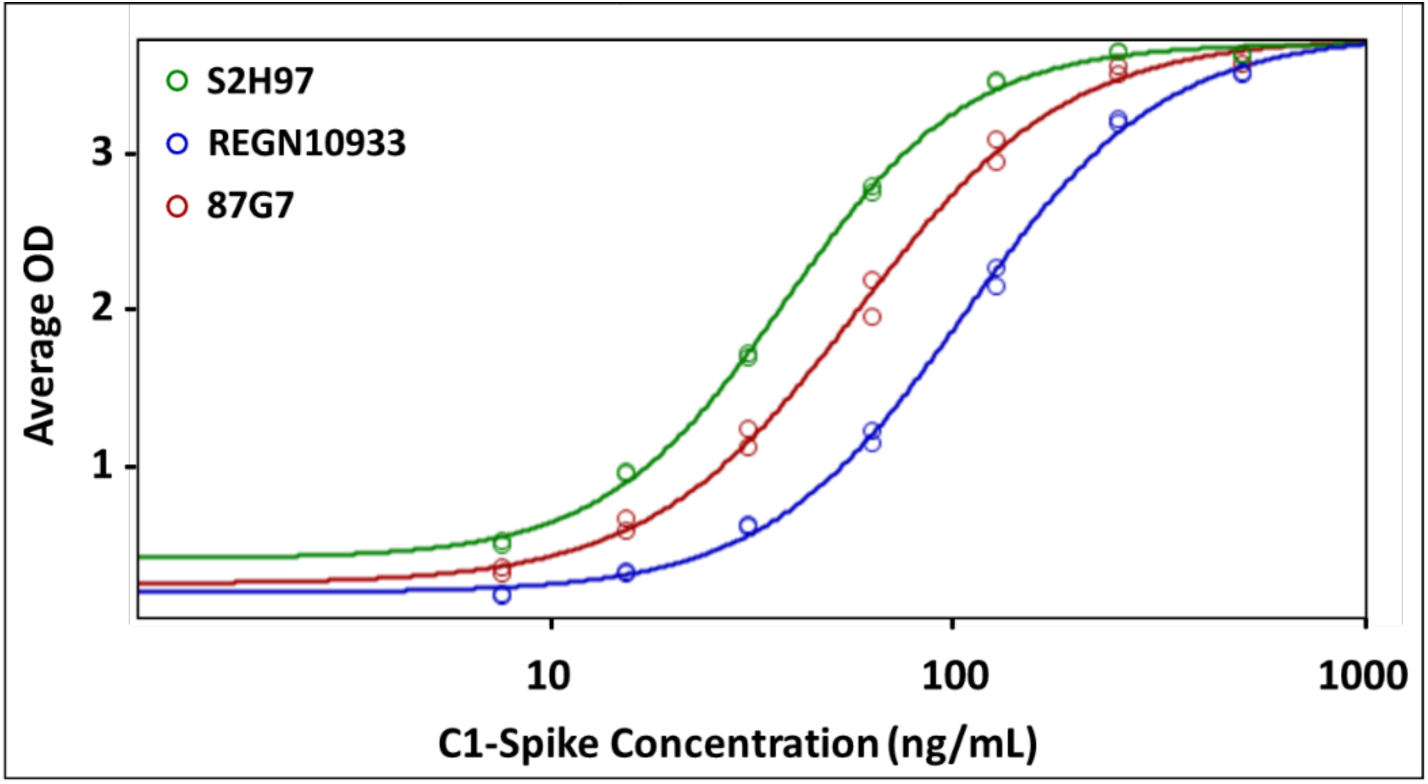
Binding curves of C1-Spike as captured by monoclonal antibodies on indirect ELISA. Binding response curves of C1-Spike concentrations to S2H97 (green), REGN10933 (blue), and in-house C1-produced 87G7 (red) antibodies.

**Supplemental Figure 4.**
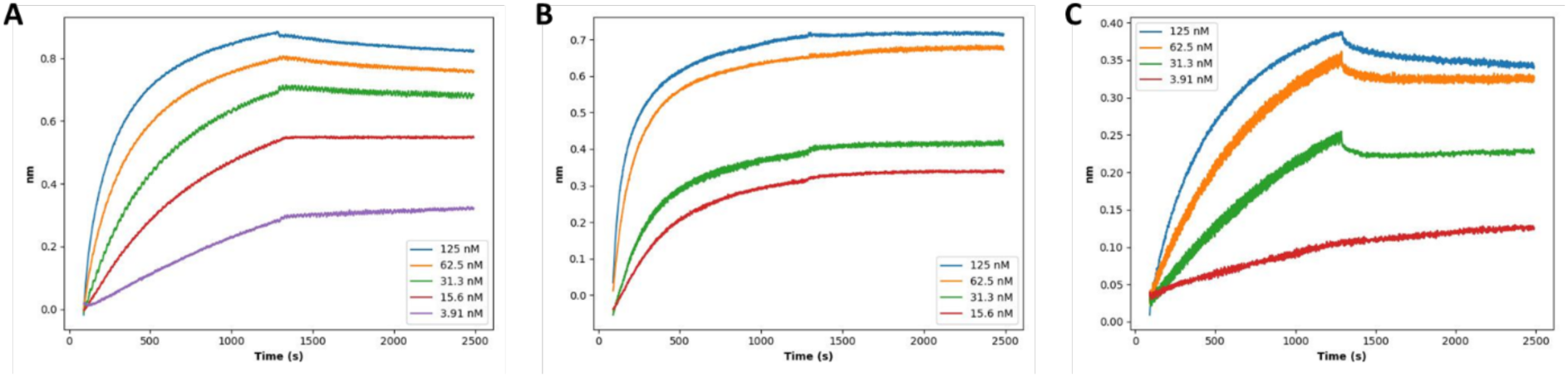
BLI curve fits for binding kinetics. (A) SARS-CoV-2 Spike binding to ACE2-Fc protein at 37°C (B) SARS-CoV-2 Spike binding to 87G7 at 37°C (C) Glycan modified SARS-CoV-2 Spike binding to ACE2-Fc protein at 37°C.

### Secondary Structure

Circular Dichroism results indicate that C1-Spike has a high percentage of α-helical structuring when compared to spike proteins expressed in other platforms (Figure 3, raw data and curve fits are found in Supplemental Figure 5). While α-helices are the primary structural component of C1-Spike, “other” structures, including 3_10_- and π-helices, bends, β bridges, and irregular or disordered structures, are the leading components of spike produced in Sf9 or CHO systems. Of note, CD spectra indicate that the six-proline stabilized HexaPro spike protein (CHO-6P-Spike) exhibits a higher ratio of alpha helical structures than its counterpart with only two proline stabilizations (CHO-Spike). The increased proline mutations were implemented to improve expression, but the protein has also displayed increased thermal stability over CHO-Spike [15]. Thus, the alpha helical structuring may be a contributor to its stability, and in turn, the high alpha helical structure detected in C1-Spike may corroborate the thermal stability of the protein later observed in Figure 4. The high beta sheet content in Sf9-Spike and CHO-Spike may additionally signal a reduction in protein stability due to limited helical stabilization when compared to C1-Spike or HexaPro [52, 53, 54].

**Figure 3.**
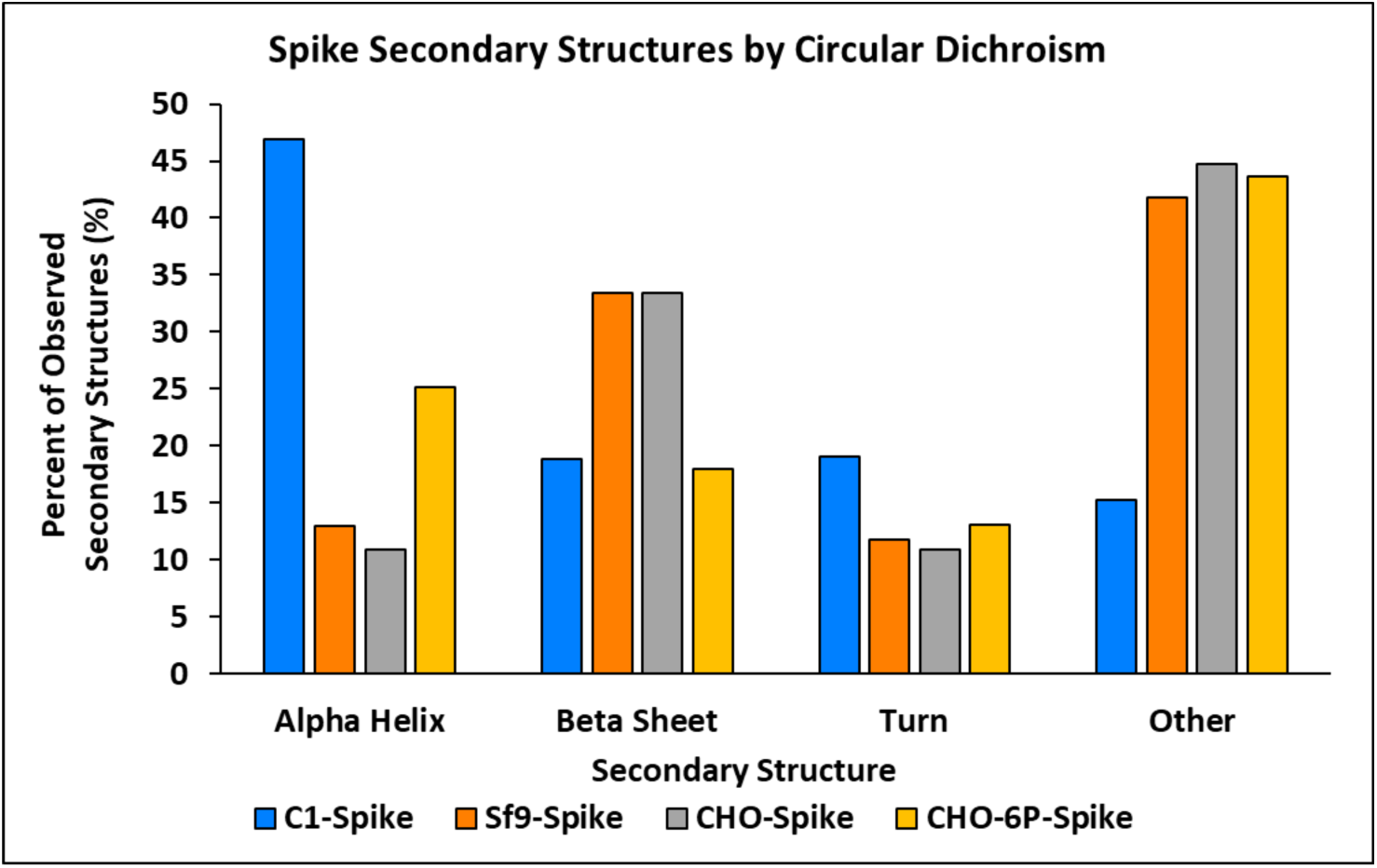
Protein secondary structural composition as probed by circular dichroism for spike proteins produced by various expression platforms. Unless otherwise noted, all proteins have 986-987 KV-PP stabilization. Percent frequency of observed structure is plotted against structure class. C1, *Thermothelomyces heterothallica* C1; Sf9, *Spodoptera frugiperda*; CHO, Chinese hamster ovary; 6P, HexaPro stabilization.

**Figure 4.**
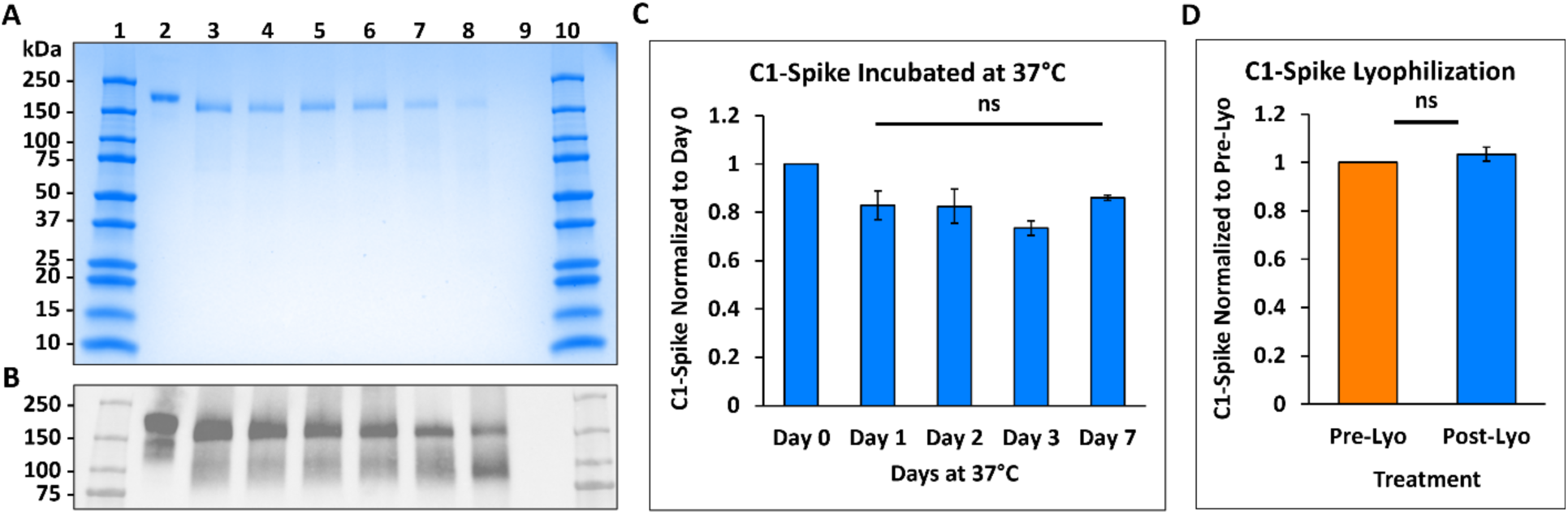
C1-Spike stability during heat or lyophilization treatments. A) SDS-PAGE stained with Coomassie Blue. Lane 1, MWM; lane 2, reference CHO-Spike (BEI NR-53937); lane 3, C1-Spike stored at -80°C; lane 4, C1-Spike stored at 4°C for 7 days; lanes 5-8, C1-Spike stored at 37°C for 1, 2, 3, and 7 days respectively; lane 9, empty; lane 10, MWM. B) Western blot for samples run in A. C) ELISA results of C1-Spike after incubation at 37°C as normalized to Day 0 sample. Error bars represent the standard deviation of triplicate measurements. One-way ANOVA of samples after incubation in 37°C determine a p-value of 0.106, showing no significant difference between time points. D) C1-Spike as measured on ELISA before and after lyophilization treatment. Error bars represent the standard deviation of triplicate measurement averages of four replicates. Student’s two-tailed t-test results in a p-value of 0.096 with no significant difference between treatments. Lyo, lyophilization; ns, not significant (p > 0.05).

**Supplemental Figure 5.**
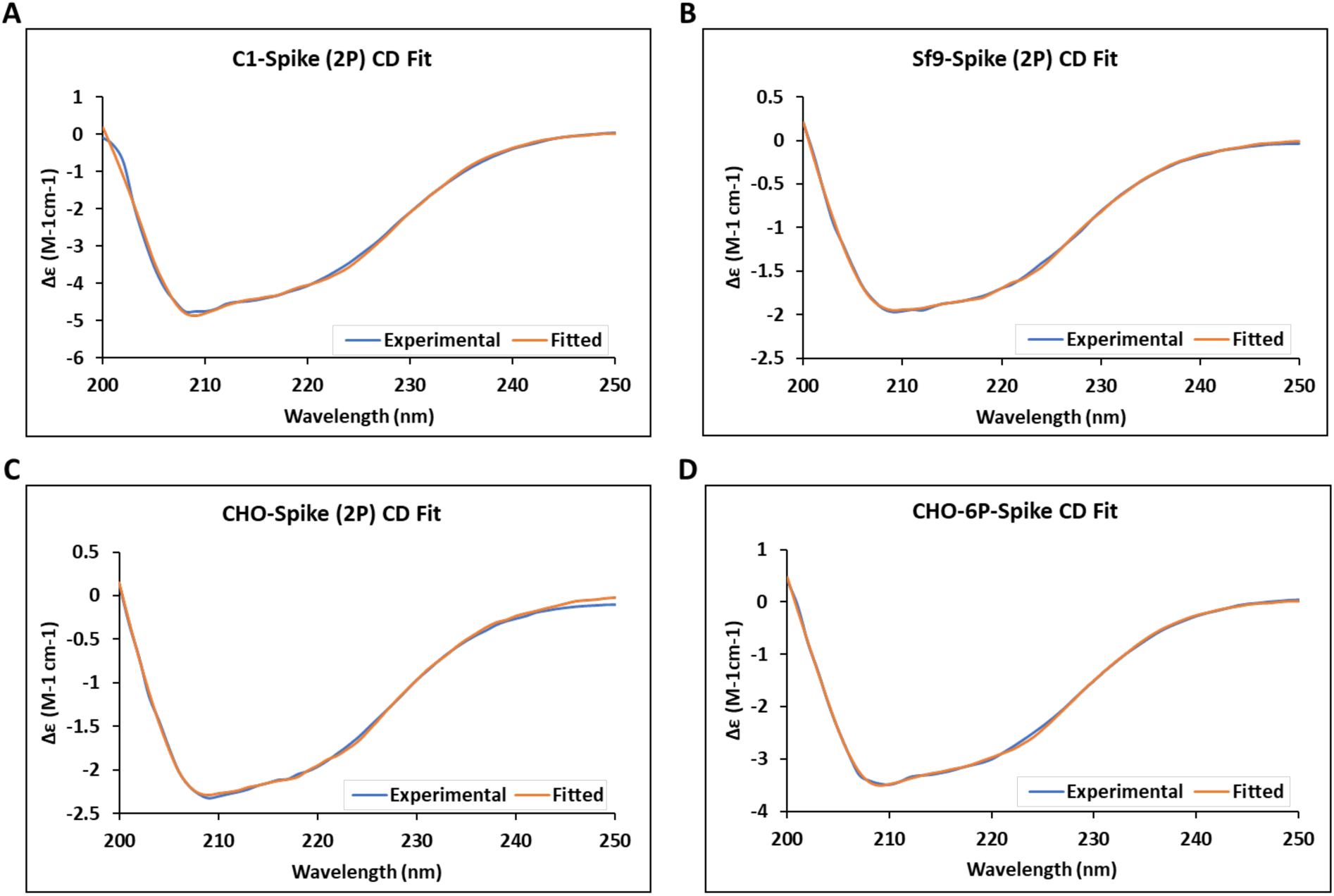
Raw spectral data of spike from circular dichroism as analyzed with BeStSel webserver. Δε (M^-1^cm^-1^) is plotted against wavelength (nm) for spike proteins produced in C1 (A), Sf9 (B), and CHO (C and D) systems. CD, circular dichroism; 2P, KV-PP proline mutation; 6P, HexaPro stabilization; C1, *Thermothelomyces heterothallica* C1; Sf9, *Spodoptera frugiperda*; CHO, Chinese hamster ovary.

### Temperature and Lyophilization Stability

Temperature stability of C1-Spike was assessed at 37°C for 1 week; SDS-PAGE and Western blot show that any degradation of the protein is predominantly one byproduct around 100 kDa (Figure 4A, B), however ELISA indicates that there is no difference in binding between samples after incubation (Figure 4C). Statistical tests determined that the difference is not significant, and any observed difference is likely explained by assay variability. Samples exposed to increased temperature show less than a 20% difference from the initial starting material, the accepted range for intra-assay validation [55, 56]. One-way ANOVA of samples after incubation in 37°C determine a *p*-value of 0.106, showing no significant difference between time points. Thus, it is likely that the shortened product of C1-Spike retains its binding epitope with 87G7 [23] as shown by ELISA and may therefore remain antigenic.

Lyophilization of C1-Spike was also considered, with no perceived impact on protein binding in ELISA (Figure 4D). Data indicate no significant difference in the sample before and after the freeze-drying treatment, where a student’s two-tailed t-test results in a *p*-value of 0.096.

### LC-MS/MS Analysis of N-Glycosylation Profile

Purified C1-Spike protein was subjected to glycoproteomic analysis before and after glycan modification reactions to characterize site-specific glycosylation and the changes introduced by glycan modification, as illustrated in Figure 5 and Supplemental Figure 6. Prior to modification reactions, the C1-Spike protein contained mostly undecorated and high-mannose glycans with compositions of HexNAc(2)Hex(4)Fuc(0)NeuAc(0) and HexNAc(2)Hex(6)Fuc(0)NeuAc(0), respectively (Figure 5A), where HexNAc is N-acetylhexosamine, Hex is hexose, Fuc is fucose and NeuAc is N-acetylneuraminic acid or sialic acid. After the glycan modification, C1-Spike protein contained mostly mono-sialylated glycans with compositions of HexNAc(3)Hex(5)Fuc(0)NeuAc(1) and HexNAc(4)Hex(5)Fuc(0)NeuAc(1) (Figure 5B). Additionally, the relative abundances of initially observed undecorated and high-mannose glycans were reduced after the glycan modification, further validating that the glycosylation pattern of C1-Spike was changed. Site-specific glycoproteomic analysis shows sialylated structures are prominent at sites N124, N167, N284, N333, N345, N659, N719, N803, N1136, N1160, N1175 after N-glycan modification. After N-glycan modification, undecorated and high-mannose structures remained prominent at certain sites, such as N19, N63 and N236.

**Figure 5.**
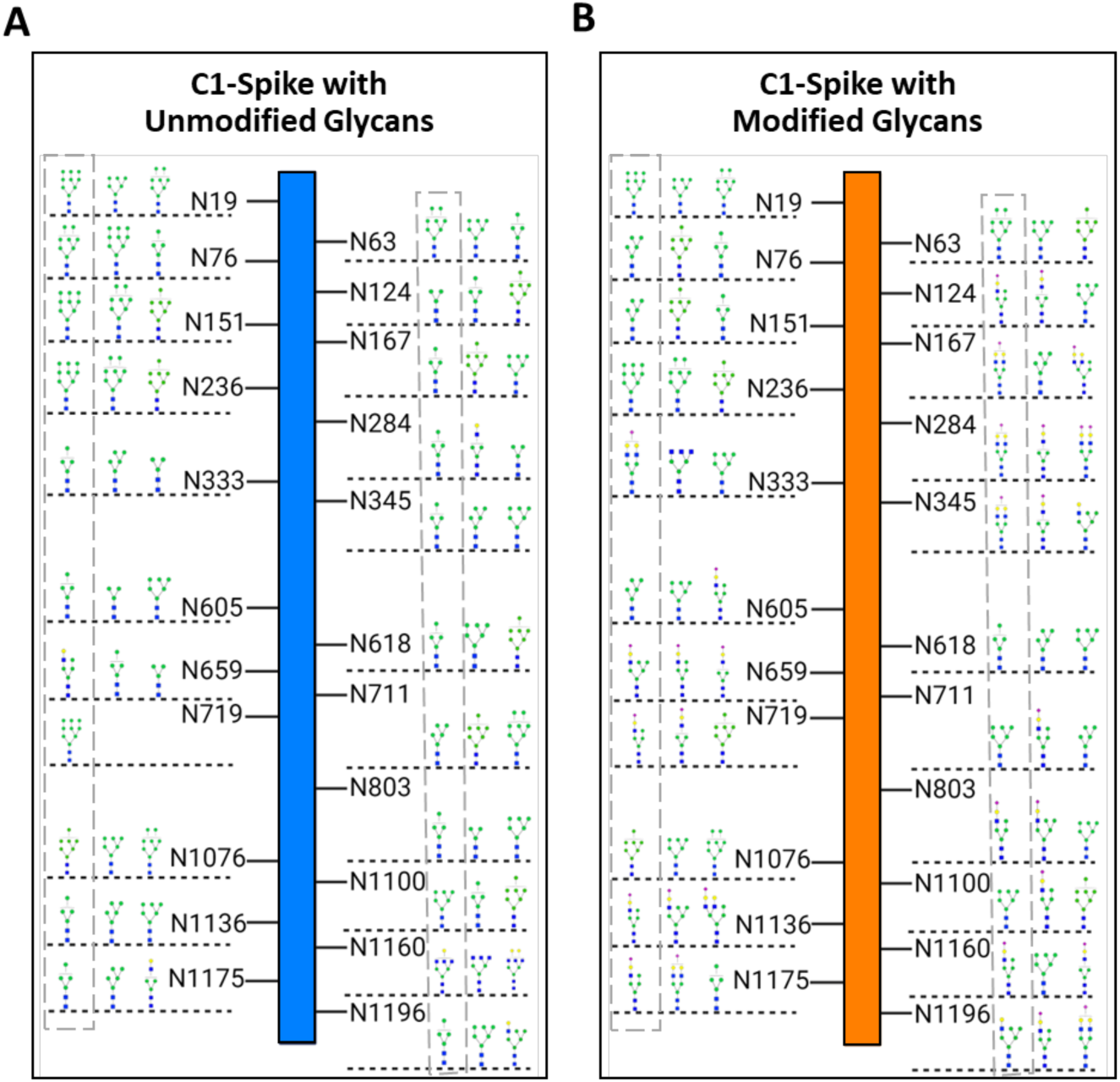
Top three most abundant N-glycan structures on each site of C1-Spike before and after glycan modification reactions. A) Mostly high-mannose and undecorated structures are observed prior to modification reactions. B) After modification, sialylated structures appear prominent at sites N124, 167, 284, 333, 345, 659, 719, 803, 1136, 1160, 1175. Dashed grey boxes represent the most abundant glycan structure at each N-linked glycosylation site.

**Supplemental Figure 6.**
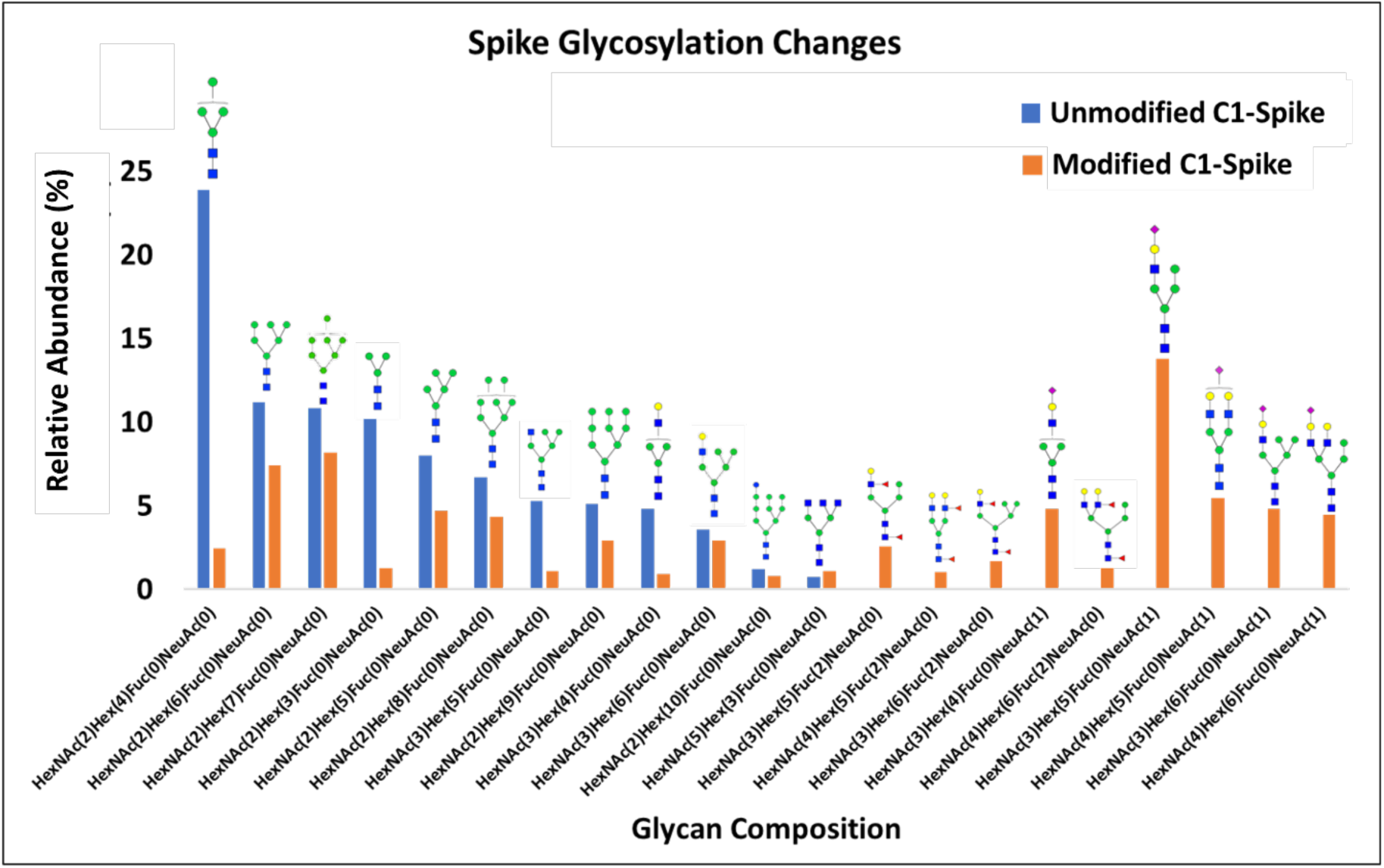
Relative abundance of N-glycans detected on C1-Spike before (blue) and after (orange) modification. N-glycans were detected on E.VFnATR.F sequence where (.) represents cleavage sites. Glycans that were detected with > 1% abundance are summarized above. The majority of glycans found on C1 Spike are undecorated N-glycan structures.

### Steered Molecular Dynamics

Supplemental Figure 7 presents a schematic of the all-atom model featuring two glycosylation patterns: high mannose (unmodified glycans) and modified glycans using the most abundant structures, as highlighted by dashed grey boxes in Figure 5. These glycosylation patterns correspond to site-specific glycan modifications, as illustrated in Figure 5, and provide a visual context for understanding the glycan modifications discussed in the following simulation results. The classification of glycans, determined by the percentage of oligomannose content present in each glycan (See Supplemental Figure 6), offers further insights into the structural variations and their potential impact on protein interactions.

This study uses SMD simulations with an all-atom model in explicit water to investigate the impact of glycan modifications on protein-protein interactions between ACE2 and the C1-Spike protein. In the simulations represented in Figure 6, the force between ACE2 and the C1-Spike trimer is plotted against the center-of-mass (COM) pulling distance. The force profiles for both unmodified (green) and modified (purple) glycan models show a sharp increase to a peak, indicating the strongest interaction, before decreasing as the COM distance extends. For the unmodified glycan case, the interaction between the unmodified C1-Spike and ACE2 complex exhibits longer-range interactions compared to the modified C1-Spike and ACE2 complex. Notably, the modified glycan model shows a higher peak force, suggesting a stronger initial interaction. These computational findings align with experimental binding data from BLI measurements (see Table 1). The K_D_ for ACE2 interaction with C1-Spike decreased from 48.8 nM (unmodified) to 2.54 nM (N-glycan modified), indicating a stronger binding affinity for the modified C1-Spike protein. This corresponds with the higher peak force observed in SMD simulations. However, binding affinity values in the literature for SARS-CoV-2-ACE2 interactions vary widely, as reported in different studies [15, 57]. Therefore, it remains unclear whether the impact of glycans on binding affinity with ACE2 is significant. Overall, glycan modification does not appear to significantly influence viral C1-Spike binding.

**Figure 6.**
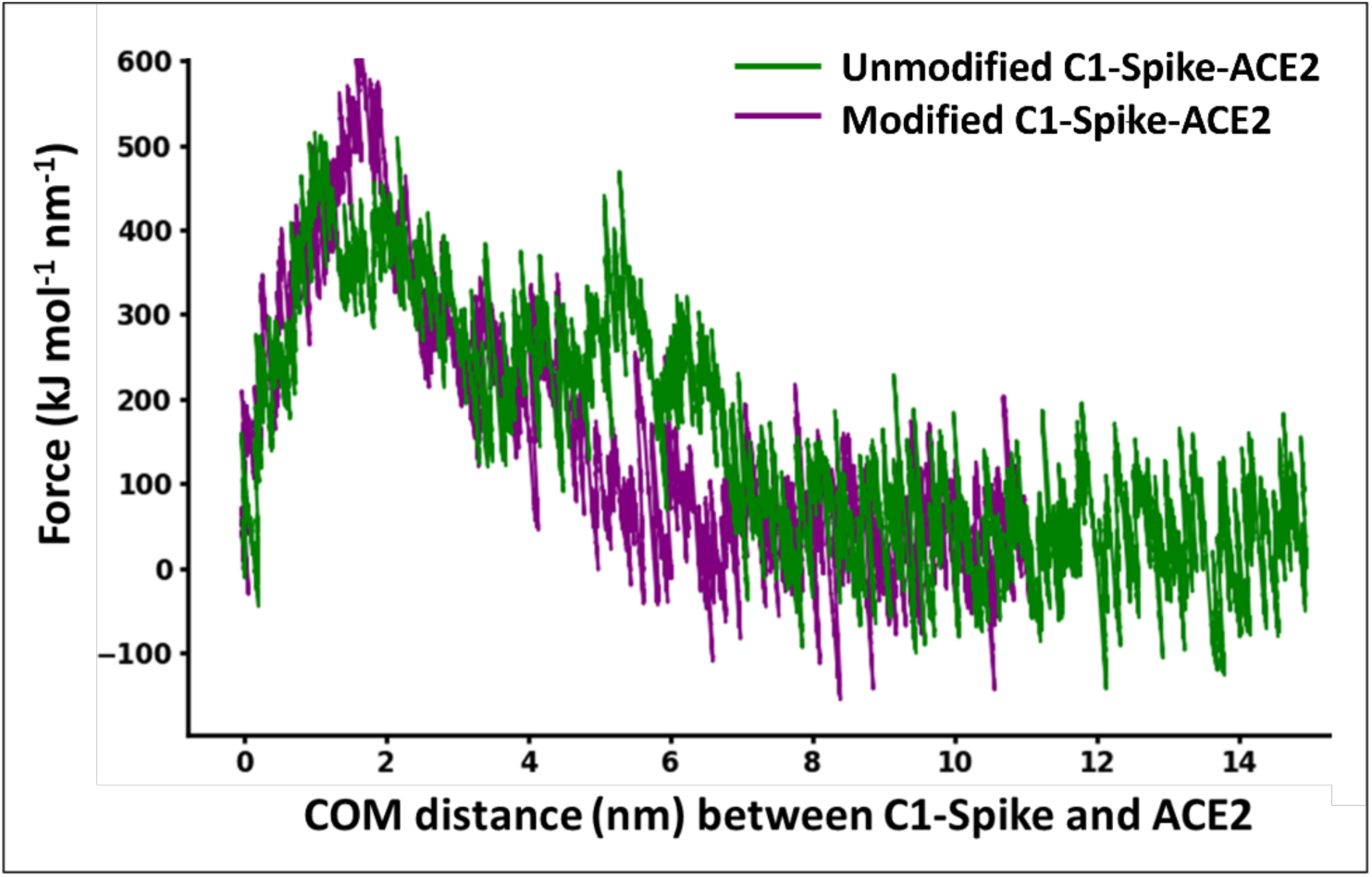
A graphical representation of the force against center of mass distance (COM) for C1-Spike interacting with the ACE2 complex. The comparison between C1-Spike with unmodified glycans (green) and modified glycans (purple) demonstrates that glycan modifications lead to a higher peak force, indicating stronger initial interactions.

**Supplemental Figure 7.**
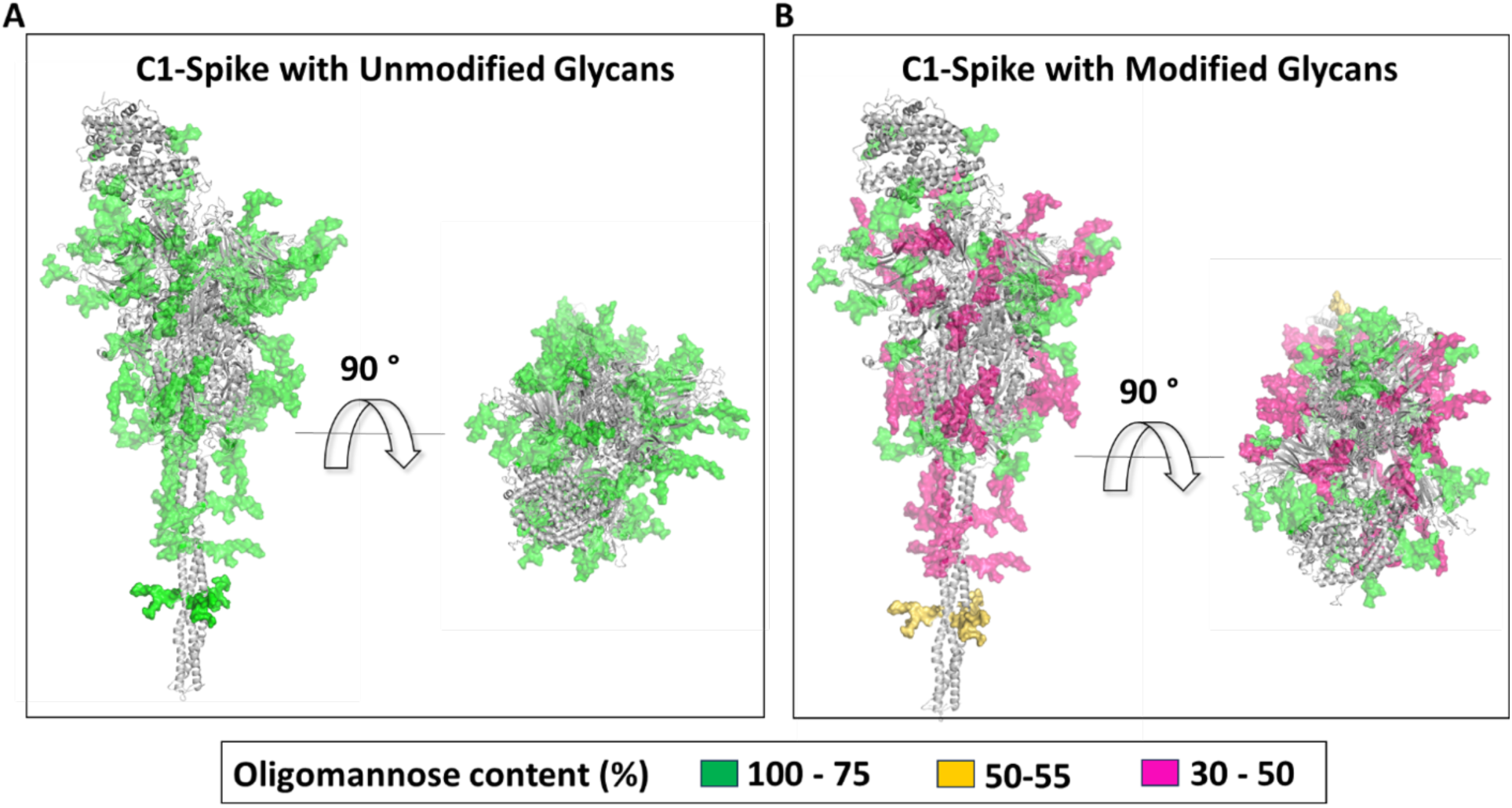
Modelled glycans used in pulling simulations. A) Most abundant glycans found on C1-Spike, B) most abundant glycans on C1 spike after

### Immunogenicity Assessment

The immunogenicity of C1-Spike in mice was compared to that of spike protein produced via the conventional HEK293 expression system. Serum spike-specific binding antibody responses and SARS-CoV-2-specific neutralization capacity of these antibodies were assessed 6 weeks after the first vaccine dose. Serum titers of spike-specific total IgG were measured by ELISA, and the virus-specific neutralization activity of these spike-specific antibodies were tested using a SARS-CoV-2 pseudovirus neutralization assay. Both C1-Spike- and HEK293-Spike-immunized mice exhibited robust serum titers of spike-specific total IgG, with no significant differences in endpoint titers between groups (Figure 7A, B). Consistent with antigen-specific binding antibody levels, both C1-Spike- and HEK293-Spike protein-vaccinated mice showed similarly robust serum-neutralizing antibody responses against SARS-CoV-2 (Figure 7C). Serum titers and neutralization capacity demonstrate the possible application of C1-Spike for use as a vaccine antigen candidate, and a competitive alternative to antigens produced in a mammalian system.

**Figure 7.**
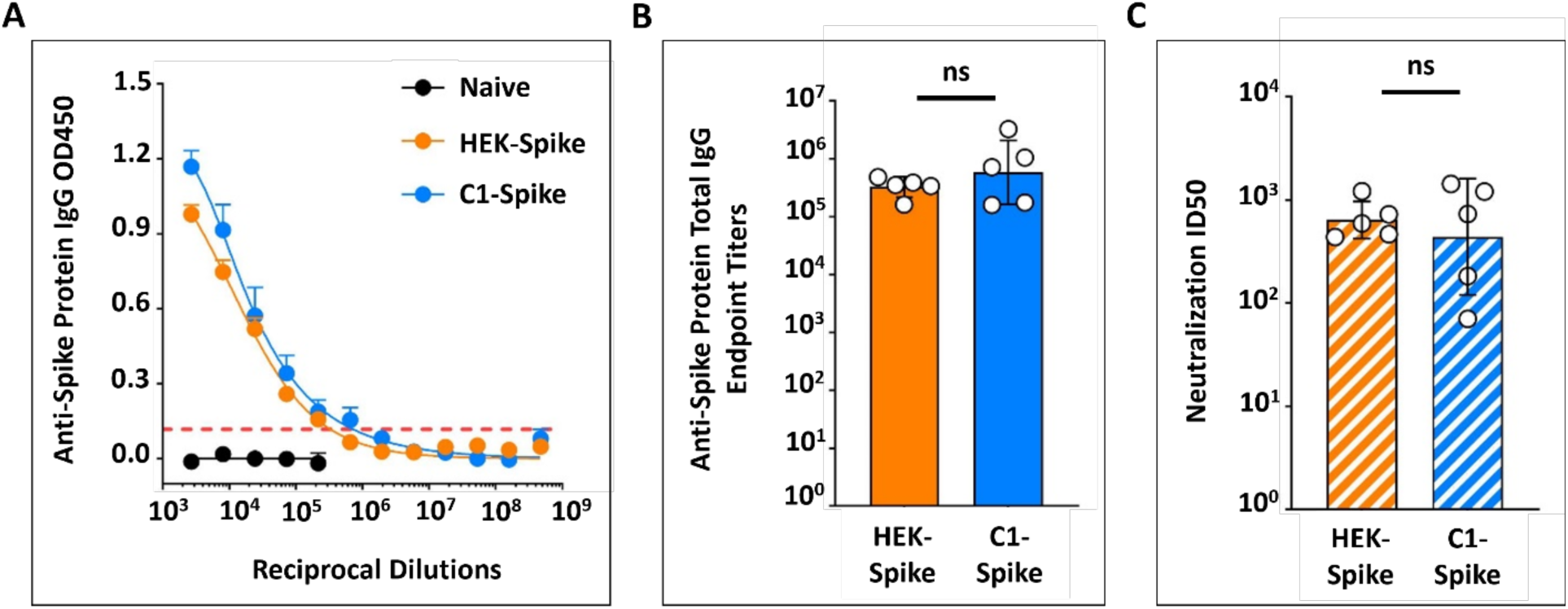
Murine humoral responses to immunization with HEK-Spike and C1-Spike. A) Anti-spike protein total IgG serum titers from mice 6 weeks after primary immunization (mean ± SEM) were measured by ELISA using HEK-Spike coated plates. B) Anti-spike protein total IgG endpoint titers (geometric mean ± SD) were calculated from titers in (A). C) Serum SARS-CoV-2 neutralizing antibody titers (ID50) (geometric mean ± SD) from mice 6 weeks after primary vaccination were measured by a pseudovirus neutralization assay. Groups in (B) and (C) were compared by independent t-test on log_10_-transformed data; ns, not significant (p > 0.05).

## Conclusions

The results presented in these studies demonstrate the ability of a *Thermothelomyces heterothallica* C1 strain expression platform to produce high-quality functional spike glycoprotein rapidly and at a high titer using low-cost media without the need for viral clearance. The strain contains a strong constitutive promoter for high expression of SARS-CoV-2 Spike protein, residues 16-1209, with mutated furin cleavage site, double proline stabilization, added foldon trimerization domain, and C-tag (Supplemental Figure 1). C1-Spike is secreted into fermentation media, allowing for easy downstream processing consisting of buffer exchange and concentration by TFF before affinity purification. Initial production processes without optimization resulted in C1-Spike titers higher than 300 mg/L in under 5 days fermentation time (volumetric productivity of 0.06 g/(L-day)), and single-step chromatography reaching nearly 90% purity (Figures 1, 2). Optimization of fermentation and purification schemes will likely increase titer and purity.

C1-Spike was thoroughly characterized to confirm size, sequence, binding capacity, and structure. Protein size, as visualized by SDS-PAGE and Western blot, shows expected molecular weight around 160 kDa, comparable to spike protein produced in CHO cells (Figures 1, 2). The protein sequence was confirmed via LC-MS (Supplemental Figure 2). Binding characteristics were shown via ELISA that C1-Spike binds similarly to 87G7, S2H97, and REGN10933 monoclonal antibodies (Supplemental Figure 3). BLI analysis also demonstrates strong binding kinetics of C1-Spike to 87G7, as well as native C1-Spike and N-glycan modified C1-Spike to ACE2, indicating functionality of C1-Spike, regardless of glycosylation profile (Table 1). These data fall within the reported range of spike antigen binding [23, 51]. Finally, the secondary structure of C1-Spike, as identified experimentally by CD, shows increased alpha helix structuring when compared to spike proteins produced in other systems (Figure 3). Of notable difference, CD results show a higher percentage of α-helical structures in C1-Spike than in HexaPro, which is in turn higher than the double proline mutants produced in CHO or Sf9. This increase in α-helix structures in C1-Spike may indicate helical stabilization of proteins produced in the C1 system and an increase in stability of the protein [52, 53, 54].

The stability of C1-Spike was investigated as it may be applicable to processing, storage, or shipping conditions. When C1-Spike is kept under 37°C conditions for one week, it is noted that there is degradation into predominantly one product, however, the minimal decrease in signal on ELISA indicates that the binding epitope is retained on the byproduct and therefore may remain antigenic (Figure 4). Stability of C1-Spike at 37°C is supported by high α-helix report in CD data (Figure 3) and allows for N-glycan modification incubation processes at 30°C overnight. C1-Spike lyophilization is also demonstrated with no impact on protein binding by ELISA (Figure 4). These data support common concerns for storage and shipping of protein subunit vaccines, as the product may be lyophilized and is likely not impacted by thermal fluctuations during these processes.

Initial glycosylation profile of C1-Spike shows heterogeneity of N-glycans, with undecorated glycans and high-mannose content (Figure 5). LC-MS/MS analysis shows successful modification of the profile, resulting in sialylated glycans (Figure 5). As a human-like glycosylation pattern is shown to impact immunogenicity [58, 59], successful N-glycan modification of C1-Spike may increase efficacy of the antigen as a vaccine candidate. BLI and SMD data indicate that N-glycan modification did not significantly affect the binding kinetics to the ACE2 receptor (Table 1 and Figure 6), however the demonstrated ability to modify the native glycan profile of C1 with a heavily glycosylated protein such as Spike establishes the ability to leverage the C1 platform to produce other proteins where the glycan profile may be critical to performance, such as therapeutic antibodies.

Finally, an initial murine immunogenicity study indicates that C1-Spike vaccinated mice produce strong serum titers of spike-specific total IgG, with no significant difference in groups when compared to mice immunized with an HEK293-produced spike antigen (Figure 7). Additionally, both groups shared similar serum-neutralizing antibody responses against SARS-CoV-2 pseudovirus (Figure 7). Collectively, the results of these pilot murine studies support the feasibility of using C1-Spike as a recombinant protein antigen in protein subunit vaccines against SARS-CoV-2.

The production of spike protein in the C1 system demonstrates the capacity of this platform to produce complex, functional, low-cost recombinant glycoproteins at flexible commercial scales. This presents C1 as a strong expression platform for recombinant protein therapies and a candidate for production of additional SARS-CoV-2 protein subunit antigens, including VOCs [60, 61], along with other respiratory viruses such as influenza [21] and Respiratory Syncytial Virus. In tandem, with the ability to modify the native N-glycan profile in a heavily glycosylated protein such as spike, the data presented establishes a potential workflow to produce other relevant human therapeutics. C1-Spike shows comparative performance in binding and structural studies, along with a possible increase in protein stability when compared to spike antigens produced in other platforms. In addition to protein performance, other advantages of this platform are the speed of production of C1, with a doubling time of 2.5 h instead of 24 h for CHO [13], high productivity using low cost standard fermentation media that does not require the typical serum or growth factors which drastically increase the cost of mammalian growth media, and a simplification of downstream processing without the need for viral clearance, saving additional time for release and cost [62]. Additionally, capacity for lyophilization and temperature stability of C1-Spike allow for a reduction in costs of shipping and storage when considering alternative vaccine platforms like mRNA.

## Acknowledgements

This work was performed under the financial assistance (award number 70NANB22H017) from U.S. Department of Commerce (DOC), National Institute of Standards and Technology (NIST) through BioMADE to UC Davis. AT and RF also acknowledge funding through the Black Family Endowment at Texas Tech University. Authors would like to acknowledge that the following reagents were obtained through BEI Resources, NIAID, NIH: Human Embryonic Kidney Cells (HEK-293T) Expressing Human Angiotensin-Converting Enzyme 2, HEK-293T-hACE2 Cell Line, NR-52511; Spike Glycoprotein (Stabilized) from SARS-Related Coronavirus 2, Wuhan-Hu-1 with C-Terminal Histidine and Twin-Strep^®^ Tags, Recombinant from CHO Cells, NR-53937; Spike Glycoprotein (Stabilized) from SARS-Related Coronavirus 2, Wuhan-Hu-1 with C-Terminal Histidine Tag, Recombinant from Baculovirus, NR-52308. The authors would like to acknowledge the Protein Structure and Dynamics Core, Biochemistry and Molecular Medicine, University of California, Davis for obtaining CD spectral data of purified proteins and the Drug Discovery Recharge Unit of UC Davis for obtaining biolayer interferometry (BLI) data. Authors would like to acknowledge that the antibodies used in ELISA binding assays, S2H97 and REGN10933 were provided by the Ilya Finkelstein lab at University of Texas at Austin, Michael Betenbaugh lab at John Hopkins University, and Steven Cramer lab at Rensselaer Polytechnic Institute.

## Data Availability Statement (DAS)

All authors agreed that in this research paper or publication all the data, methodology, and supporting findings can be accessed publicly and we do not have any restrictions on access.

